# Modulation of stress-related behaviour by hypothalamic engagement of preproglucagon neurons in the nucleus of the solitary tract

**DOI:** 10.1101/2022.02.04.479117

**Authors:** Marie K. Holt, Natalia Valderrama, Maria J Polanco, Linda Rinaman

## Abstract

Stress-induced behaviours are driven by complex neural circuits and some neuronal populations concurrently modulate diverse behavioural and physiological responses to stress. Glucagon-like peptide-1 (GLP1)-producing preproglucagon (PPG) neurons within the lower brainstem caudal nucleus of the solitary tract (cNTS) are particularly sensitive to stressful stimuli and are implicated in multiple physiological and behavioural responses to interoceptive and psychogenic threats. However, the afferent inputs driving stress-induced activation of PPG neurons are largely unknown, and the role of PPG neurons in anxiety-like behaviour is controversial. Through chemogenetic manipulations we reveal that cNTS PPG neurons have the ability to moderately increase anxiety-like behaviours in mice in a sex-dependent manner. Using an intersectional approach, we show that a corticotropin-releasing hormone (CRH)-rich input from the paraventricular nucleus of the hypothalamus (PVN) drives activation of both the cNTS as a whole and PPG neurons in particular in response to acute stress. Finally, we demonstrate that NTS-projecting PVN neurons are necessary for the ability of acute stress to suppress food intake. Our findings reveal sex differences in behavioural responses to PPG neural activation and highlight a hypothalamic-brainstem pathway in stress-induced hypophagia.

## 1. Introduction

Behavioural and physiological responses to stressors are essential for survival and are tightly controlled by the brain. Exposure to stressful stimuli leads to activation of the hypothalamic-pituitary-adrenal (HPA) axis, sympathetic arousal, and elicitation of adaptive behaviours, including heightened vigilance and decreased exploration and food intake (Holt & Trapp, 2016; Myers et al., 2017; Ulrich-Lai & Herman, 2009). While hypothalamic areas are often considered the primary site of stress processing and modulation (Bains et al., 2015), multiple brainstem regions are essential for the integration of signals of both interoceptive sensory and psychogenic stress (Itoi & Sugimoto, 2010; Myers et al., 2017). The caudal part of the nucleus of the solitary tract (cNTS) is particularly well-positioned to integrate such signals (Holt, 2021; Maniscalco & Rinaman, 2017, 2018). Neurons within the cNTS are sensitive to both interoceptive and psychogenic stressors (Maniscalco & Rinaman, 2017), presumably mediated by their receipt of interoceptive sensory input from the spinal cord and vagal afferents along with descending input from brainstem regions, the cerebral cortex, limbic system, and hypothalamic nuclei, including the paraventricular nucleus of the hypothalamus (PVN) (Holt, Pomeranz, et al., 2019; Leon et al., 2021; Rinaman & Rothe, 2002; van der Kooy et al., 1984).

Psychogenic stressors elicit robust suppression in feeding which can last hours to days in rats (Calvez et al., 2011; Vallès et al., 2000). Circuits driving this stress-induced hypophagia partially overlap with satiation circuits (Calvez et al., 2011) that include neurons within the cNTS. Multiple cell types within the cNTS have the ability to modulate feeding (D’Agostino et al., 2016, 2018; Roman et al., 2016, 2017; Wang et al., 2015), including glucagon-like peptide-1 (GLP1)-expressing PPG neurons. Indeed, in mice, chemogenetic and optogenetic activation of cNTS PPG neurons robustly suppresses food intake (Brierley et al., 2021; Cheng et al., 2020; Gaykema et al., 2017; Holt, Richards, et al., 2019; Liu et al., 2017) and cNTS PPG neurons are necessary for limiting intake of large meals (Brierley et al., 2021; Holt, Richards, et al., 2019).

However, interfering with endogenous, central GLP1 signalling either pharmacologically, genetically, or through inhibition of PPG neuron activity has limited impact on *ad libitum* feeding in mice (Brierley et al., 2021; Cheng et al., 2020; Ghosal et al., 2017; Holt, Richards, et al., 2019; Liu et al., 2017; Terrill et al., 2019; Williams et al., 2018). Evidence suggests that rather than driving meal-induced satiation under normal physiological conditions, PPG neurons are more strongly engaged by stressful conditions that inhibit food intake. PPG neurons are activated by a variety of acute stressors in both rats and mice (Leon et al., 2021; Maniscalco et al., 2015; Rinaman, 1999; Terrill et al., 2019), and GLP1 acts centrally to generate stressor-like physiological responses that include activation of the HPA axis (Gil-Lozano et al., 2010; Kinzig et al., 2003) and increased heart rate (Barragán et al., 1999; Holt et al., 2020). Additional evidence indicates that PPG neurons are necessary for stress-induced hypophagia in mice (Holt, Richards, et al., 2019), that stress-induced hypophagia is significantly reduced in rats after central blockade of GLP1 receptors (Maniscalco et al., 2015; Zheng et al., 2019), and that central endogenous GLP1 elicits anxiety-like behaviours in rats (Kinzig et al., 2003; López-Ferreras et al., 2020; Maniscalco et al., 2015; Zheng et al., 2019). Surprisingly, however, a previous study using chemogenetic activation of PPG neurons in mice failed to detect any changes in anxiety-like behaviour assessed in the elevated plus maze or open field in male mice (Gaykema et al., 2017).

The current study was designed to address two objectives: first, to reassess the role of PPG neurons in anxiety-like behaviours in male and female mice tested in the open field and to additionally test the role of these neurons in acoustic startle responses, and second, to delineate inputs from the PVN and other brain regions that potentially drive stress-induced activation of PPG neurons. We previously reported that cNTS-projecting PVN neurons are activated in response to acute restraint stress in mice (Holt, Pomeranz, et al., 2019), but the role of this projection in activating PPG neurons and driving behavioural responses to stress has not been tested. The present study used a chemogenetic approach to activate PPG neurons within the cNTS in mice, revealing that their activation promotes both hypophagia and anxiety-like behaviours, with the latter expressed in a sex-dependent manner. Using rabies-mediated monosynaptic circuit tracing and intersectional chemogenetic inhibition of neural activity, we also identify a descending pathway from the PVN to the NTS that mediates the ability of acute psychogenic stress to activate PPG neurons and suppress food intake.

## 2. Materials and Methods

Experimental protocols were approved by the Florida State University (FSU) Institutional Animal Care and Use Committee, and were consistent with the US Public Health Service’s Policy on the Humane Care and Use of Laboratory Animals and the NIH Guide for the Care and Use of Laboratory Animals.

### 2.1 Animals

Male and female mGlu-Cre/tdRFP transgenic mice were bred in house (n=38); wildtype, male C57BL/6J (stock #000664) mice (n=13) were obtained from The Jackson Laboratories. Details regarding viral injections, sample sizes, and animal ages are provided in Table 1.

**Table 1.** Experimental details

| **Experiment** | **Mouse strain** | **Viral transduction,**<br>**sample size and sex** | **Age at viral**<br>**injection**<br>**surgery**<br>**(months±SD)** | **Age at**<br>**perfusion**<br>**(months±SD)** |
| --- | --- | --- | --- | --- |
| 1) Chemogenetic<br>activation, PPG neurons | Glu-Cre/tdRFP | hM3Dq (N=8F+6M)<br>mCherry (N=8F+6M) | 3.0±1.4 | 8.87±2.2 |
| 2) Monosynaptic<br>retrograde tracing | Glu-Cre/tdRFP | RABV (N=6F,4M) | 3.5±0.12 | 3.7±0.12 |
| 3) Chemogenetic<br>inhibition, NTS-projecting<br>PVN neurons | C57BL/6J | hM4Di (N=5M)<br>GFP (N=8M) | 2.9±0.05 | 5.4±0.03 |

Mice were individually housed from the start of behavioural experiments and were kept on a 12 h light/dark cycle. They had *ad libitum* access to water and rodent chow (Purina) unless otherwise stated and were habituated to handling and injection procedures every day for at least one week prior to experimentation. mGlu-Cre transgenic mice express Cre recombinase under the control of the glucagon promoter, allowing selective targeting of GLP-1-expressing PPG neurons (Anesten et al., 2016; Holt, Richards, et al., 2019; Parker et al., 2012). These cells also express the fluorescent reporter, tdRFP, in a Cre-conditional manner (Luche et al., 2007). Local colonies of mGlu-Cre/tdRFP transgenic mice were established at FSU in 2013 from a strain received from Frank Reimann at Cambridge University (UK). The original Cambridge mGlu-Cre mice were generated in 2008 and maintained for >20 generations before receipt by FSU. At FSU, mGlu-Cre mice have been maintained for >15 generations on a C57BL/6 background.

### 2.2 Exp 1: chemogenetic activation of PPG neurons

#### 2.2.1 Stereotaxic injections

mGlu-Cre/tdRFP mice were anaesthetized using isoflurane (1-3%, 1.5ml/min in O_2_) and placed in a stereotaxic frame with the nose pointing downwards to expose the dorsal surface of the neck and facilitate access to the caudal brainstem. An incision was made through the skin along the midline extending from the occipital crest to the first vertebra and the underlying muscles were separated to expose the roof of the fourth ventricle caudal to the cerebellum. The meningeal layer was penetrated using a 30g needle and obex was visualized. To target the cNTS with AAVs (Table 2), the tip of a glass needle was inserted 400µm lateral and 100µm rostral to obex, and then lowered 350µm below the dorsal surface of the brainstem.

**Table 2.** Viral titers, injection volumes, and sources.

| **Virus** | **Titer**<br>**(pfu/ml)** | **Volume** | **Source** | **Reference** |
| --- | --- | --- | --- | --- |
| AAV8-hSyn-DIO-mCherry | $2.1 \times 10^{12}$ | 250 nl | Addgene #44361-AAV8 | (Krashes et al., 2011) |
| AAV8-hSyn-DIO-hM3Dq:mCherry | $2.2 \times 10^{12}$ | 250 nl | Addgene #50459-AAV8 | (Armbruster et al., 2007) |
| AAV8-hSyn-DIO-hM4Di:mCherry | $2.0 \times 10^{12}$ | 400 nl (PVN) | Addgene #44362-AAV8 | (Krashes et al., 2011) |
| AAV1-CAG-FLEX-EGFP | $2.0 \times 10^{12}$ | 400 nl | Addgene #51502-AAV1 | (Oh et al., 2014) |
| AAVrg-hSyn-Cre | $1.2 \times 10^{12}$ | 200 nl | Addgene #105553-AAVrg | Gift from James M. Wilson |
| AAV5-EF1a-FLEX-TVA:mCherry | $2.13 \times 10^{12}$ | 100 nl | GVC-AAV-67, Stanford Gene Vector and Virus Core | (Watabe-Uchida et al., 2012) |
| AAV8/733-CAG-FLEX-RabiesG | $2.4 \times 10^{13}$ | 100 nl | GVC-AAV-59, Stanford Gene Vector and Virus Core | (Watabe-Uchida et al., 2012) |
| (EnvA)-RABV-ΔG-GFP | $2 \times 10^8$ | 400 nl | Kevin Beier, University of California at Irvine, CA | (Wickersham et al., 2007) |

#### 2.2.2 Food intake measurements

A minimum of two weeks after stereotaxic injection, mice were individually housed in cages fitted with the BioDAQ food-intake monitoring system (Research Diets, New Brunswick, New Jersey) and left to habituate to the feeding system and handling for at least one week prior to experimentation. On test days, chow access gates were closed to prevent feeding for three hours prior to dark onset. Using a mixed-model design with a minimum of 48 hours between conditions, mice were injected 30mins before dark onset with either saline vehicle (control) or vehicle containing clozapine-*N-oxide* dihydrochloride [CNO; Tocris; 2 mg/kg, 2 ml/kg, i.p.; dose based on previous studies (Brierley et al., 2021; Gaykema et al., 2017; Holt et al., 2020; Holt, Richards, et al., 2019)]. At dark onset, food gates were opened to provide chow access, with intake measured continuously by the BioDAQ. Meal-pattern data were subsequently extracted using the BioDAQ Data Viewer (Research Diets), with a meal defined as a feeding episode in which at least 0.02g of chow was consumed with an intermeal interval of at least 300s. Cumulative food intake, average meal size, and average meal duration were analysed during the first six hours after dark onset, as well as latency to begin the first meal and first meal size.

#### 2.2.3 Acoustic startle response

Using a mixed-model design with virus as a between-subjects factor and treatment (CNO vs saline) as within-subjects factor, mice were tested in the SR-Lab-Startle Response System (San Diego Instruments, San Diego, CA) during the light phase of the photoperiod (2-10 hours after light onset). Each mouse was tested under both CNO and saline conditions using a randomised, counterbalanced design with at least 48 hours between tests. On each day of testing, mice were moved to the test room and left to acclimatize for 30 mins after which they were injected with CNO (2 mg/kg, 2 ml/kg) or saline (2 ml/kg). Thirty mins after injection, mice were placed individually into a clear acrylic enclosure which allowed mice to turn around without constraint (San Diego Instruments, San Diego, CA). Mice were left to acclimatize for 5 mins in the darkened startle chamber with a constant background noise level of 50 db. Following this initial period, mice were exposed to a series of 50 ms white noise bursts at 75, 90, and 105 db (10 repeats of each) in a randomly generated order with random intervals of 20-40 sec between noise bursts. Testing lasted 23 mins in total. All testing was done during the light cycle.

#### 2.2.4 Open field test

Using a between-subjects design (with virus as the between-subjects factor), mice were injected with CNO (2 mg/kg, 2 ml/kg) 30mins prior to individual testing in a novel open field (50cm x 50cm), with exploratory behaviour recorded for 20mins during the light phase (5-9 hours after light onset) of the photoperiod. Location was tracked using the ezTrack Location Tracker open-source software (Pennington et al., 2019). Total distance travelled and time spent in the central region (30 x 30cm) of the open field were analysed.

#### 2.2.6 Transcardial perfusion and tissue preparation

Mice were injected with either saline (2 ml/kg) or CNO (2 mg/kg, 2ml/kg) and anaesthetized using pentobarbital sodium (Fatal Plus, 100mg/kg, i.p.; Henry Schein) 90mins later. They were then transcardially perfused with ice-cold phosphate buffer (PB, 0.1M, pH 7.2) followed by 4% paraformaldehyde in PB before brains were extracted for further processing (see sections 2.5-2.6).

### 2.3 Exp 2: Rabies virus-mediated circuit tracing

Using the surgical approach described above (section 2.2.1), mGlu-Cre/tdRFP mice (n=4M, 4F) received cNTS microinjection of two helper AAVs encoding rabies glycoprotein and TVA receptor (Table 2). Three weeks later, the same mice received cNTS-targeted microinjection of EnvA-pseudotyped G-deleted rabies [(EnvA)-RABV-ΔG-GFP; Table 2]. As previously described (Holt, Pomeranz, et al., 2019), (EnvA)-RABV-ΔG-GFP was injected at each of two medial-lateral injection sites relative to obex: (1) 250µm lateral, 100µm rostral, and 450-350µm below the surface of the brainstem; and (2) 400µm lateral, 100µm rostral, and 450-350µm below the surface of the brainstem. Viral titers, injection volumes, and sources are listed in Table 2.

One week after injection of rabies virus, mice were restrained in a decapicone for 30mins or left undisturbed in their home cages. Ninety minutes after the onset of restraint stress, mice were anaesthetized and transcardially perfused as described above (section 2.2.6). Brains were extracted and processed as described in sections 2.6-2.7.

### 2.4 Exp 3: Chemogenetic inhibition of NTS-projecting PVN neurons

#### 2.4.1 Stereotaxic injections targeting cNTS and PVN

Wildtype, male C57BL/6J mice were anaesthetised using isoflurane and placed in a stereotaxic frame. The cNTS was targeted as described above (section 2.2.1) for bilateral microinjection of AAVrg-hSyn-Cre (Table 2) to induce Cre expression in cNTS-projecting neurons. Following cNTS injection in the same surgical session, the skull was levelled, and a skin incision was made along the midline of the skull overlying the diencephalon. A single hole was drilled in the skull to allow bilateral targeting of the PVN with AAV8-hSyn-DIO-hM4Di:mCherry or AAV1-CAG-FLEX-EGFP (Table 2) at the following coordinates from bregma: 820µm caudal, 100µm lateral, and 4.75mm ventral.

#### 2.4.2 Food intake experiments

To assess the effect of chemogenetic inhibition of cNTS-projecting PVN neurons on baseline food intake, chow access was prevented for three hours prior to dark onset, and CNO (2mg/kg, 5ml/kg) or saline (5ml/kg) was injected 30mins prior to dark onset. The BioDAQ system was unavailable for this experiment. Instead, home cage food intake was assessed manually 1, 2, and 4h after pre-weighed chow was returned to the hopper at dark onset.

At least 21 days later, stress-induced hypophagia was assessed in the same mice using a mixed-model design with each mouse exposed to restraint stress only once. Chow access was removed three hours prior to dark onset and all mice were injected with CNO (2mg/kg, 5ml/kg) one hour before dark onset. Beginning 30min before dark onset, mice were restrained in decapicones (MDC-200, Braintree Scientific) or left undisturbed for 30 min. Pre-weighed chow was returned to the hopper at dark onset when restrained mice were released from restraint, and cumulative food intake was manually measured 2h later.

#### 2.4.3 Transcardial perfusion

At least 12 days after assessment of food intake, the same mice were injected with CNO (2mg/kg, 5ml/kg) 1.5-3hs into the light phase. Thirty minutes later they were exposed to one of two psychogenic stressors: 30 mins restraint stress as above (n=5), or 20 mins in a novel, brightly illuminated open field (50 x 50cm). Mice were then returned to their home cage. Ninety minutes after stressor onset, mice were anesthetized and transcardially perfused with fixative as described above (section 2.2.6).

### 2.5 Immunohistochemical (IHC) and immunofluorescence labelling

After perfusion fixation, brains were immediately extracted and post-fixed overnight in 4% paraformaldehyde at 4°C. Following cryoprotection in sucrose (20% in 0.1 M PB), coronal sections (30-35µm) were collected on a freezing microtome and stored in cryopreservant solution (Watson et al., 1986) at −20°C until further processing.

Sections were removed from cryoprotectant and rinsed in four changes of 0.1M PB, followed by treatment with 0.5% sodium borohydride for 20 mins at room temperature. Sections destined for IHC (i.e., immunoperoxidase) labelling were further incubated in 0.15% hydrogen peroxide for 15 mins at room temperature to suppress endogenous peroxidase activity. Pre-treated sections were then incubated overnight at room temperature (16-24h) in one of several primary antibodies (see Table 3) diluted in 0.1M PB containing 0.3% Triton-X and 1% normal donkey serum. For anti-GLP1 labelling, sections were instead incubated for one hour at room temperature followed by 60-65h at 4°C. Sections were then rinsed in four changes of 0.1M PB over one hour followed by incubation in species-specific secondary antibody (Table 3, 1:500) at room temperature for one hour (for IHC) or two hours (for immunofluorescence). After three rinses in 0.1M PB sections were either mounted onto glass slides (for immunofluorescence) or incubated in Vectastain Elite avidin-biotin complex kit reagents diluted in 0.1M PB containing 0.3% Triton for 1.5-2h (for IHC). Peroxidase activity was localized using diaminobenzidine catalysed with hydrogen peroxide. Sections were then rinsed, mounted and left to dry. Following dehydration in increasing concentrations of ethanol, sections were cleared in xylene and coverslipped using Cytoseal 60 (Electron Microscopy Sciences, 18007).

**Table 3.** Primary antibody details.

| Target | Source and catalogue # | Antibody accession # | Dilution | Secondary antibody |
| --- | --- | --- | --- | --- |
| cFOS | Cell Signaling Technology, 9F6 | AB_2247211 | 1:10,000<br>(IHC), 1:1000<br>(IF) | Biotin-conjugated anti-rabbit<br>(IHC), AlexaFluor-647 (IF) |
| mCherry | Takara Bio, #632543 | AB_2307319 | 1:2,000 | AlexaFluor-488 anti-mouse |
| dsRed | Takara Bio, #632496 | AB_10013483 | 1:20,000<br>(IHC),<br>1:2,000 (IF) | Biotin-conjugated anti-rabbit<br>(IHC), AlexaFluor-Cy3 (IF) |
| GFP | Abcam, Ab13790 | AB_300798 | 1:5,000 | AlexaFluor-488 anti-chicken |
| GLP1 | Bachem, T-4363 | AB_518978 | 1:10,000 | Biotin-conjugated anti-rabbit |

### 2.6 RNAscope in situ hybridization

Tissue sections were processed using fluorescence *in situ* hybridisation (RNAscope Multiplex Fluorescent Reagent Kit version 2, Advanced Cell Diagnostics [ACD], #323100) to label preproglucagon (*Ppg*) mRNA transcripts in the cNTS and *Crh* mRNA transcripts in the PVN. Coronal sections containing the PVN or cNTS were pre-treated with hydrogen peroxide for 30mins (ACD, #323100) at room temperature, slide-mounted in dH_2_O (Fisherbrand SuperFrost Plus #12-550-15) and left to dry overnight at room temperature. Following 10s dehydration in 100% ethanol, a hydrophobic barrier was drawn, and sections were treated with proteinase IV (ACD, #322336) for 25mins at room temperature. Following three rinses in dH_2_O, sections were left to incubate in RNAscope probe Mm-*Ppg* (ACDbio #482311, NM_008100.4) or Mm-*Crh* (ACDbio # 316091, NM_205769.2) for 2h at 40°C in the HybEZTM oven (ACD). Sections then underwent three amplification steps and labelling with Cy3- or Cy5-conjugated Tyramine Signal Amplification Plus (PerkinElmer) according to the ACD protocol. Slides were washed in wash buffer (ACD, #310091) 3 x 3 mins between incubation steps. The same sections were subsequently processed as detailed above for immunofluorescent enhancement of GFP or tdRFP/mCherry reporter proteins.

### 2.7 Light microscopy and cell counting

Images of single or dual IHC/immunofluorescence labelling were captured using a KEYENCE microscope (BZ-X700) and integrated software to generate focused images though the section thickness. Fluorescence immunolabelling and *in situ* hybridization were visualized on a Leica TCS SP8 confocal microscope using a 20x air objective and a 40x oil-immersion objective. AlexaFluor-488, Cy3, and AlexaFluor-647 were excited using a 488nM OPSL, 552nM OPSL, and 638nm Diode laser, respectively. Confocal images were acquired sequentially using Leica LAS 4.0 image collection software. Brightness and contrast were adjusted using Fiji open source biological image analysis software (Schindelin et al., 2012). All cell counts were performed manually in captured images. In Exp 3, counts of cFOS-IR nuclei in the cNTS (approximately 7.5-8.0 mm caudal to bregma) were made using an average of 5 sections per mouse, with sections spaced by 105µm.

### 2.8 Statistics

Statistical significance was assessed using null-hypothesis testing, including Student’s T test and 3- and 2-way ANOVA, as indicated in the text and figures. Statistically significant interactions (p<0.05) were followed up with Sidak’s multiple comparisons test. Exact p-values for each test are indicated in graphs and/or figure legends. In each graph, individual data points are indicated either by a symbol (if unpaired) or by a line (if paired). To facilitate interpretation of the magnitude and precision of the results, we also include plots of relevant effect sizes derived from estimation statistics, in which the mean difference between groups is plotted on a floating axis as a bootstrap sampling distribution adjacent to each traditional bar graph. The mean difference is depicted as a cross (x); the 95% confidence interval is indicated by the vertical error bar. Effect size plots were generated using estimationstats.com (Ho et al., 2019).

## Results

### Selective chemogenetic activation of PPG neurons potently suppresses feeding by driving meal termination

We first verified the selectivity of the transgenic mGlu-Cre/tdRFP mouse model using RNAscope *in situ* hybridisation to detect *Ppg* mRNA. Cre-conditional fluorescent reporter tdRFP was fully colocalised with *Ppg* mRNA expression within the cNTS (Fig S1A). Injection of AAV8-DIO-hM3Dq:mCherry or AAV8-DIO-mCherry into the cNTS (Fig 1A) led to high transduction efficiency and selectivity (Fig 1B,C) indicated by colocalization of mCherry (viral transgene) and *Ppg* mRNA (Fig 1B, C), with no difference in transduction efficiency between the two viruses (Fig S1B). Compared to saline injection, CNO (2 mg/kg, i.p.) increased the percentage of cFOS-positive PPG neurons in the cNTS of hM3Dq-expressing mice by 52 percentage points (95% confidence interval [CI]: 41.9 to 61.5 percentage points; Fig 1D,E). This CNO effect was independent of sex (Fig S1C) and was absent in virus control mice (95%CI: −4.05 to 4.95 percentage points; Fig 1D,E).

**Figure 1.**
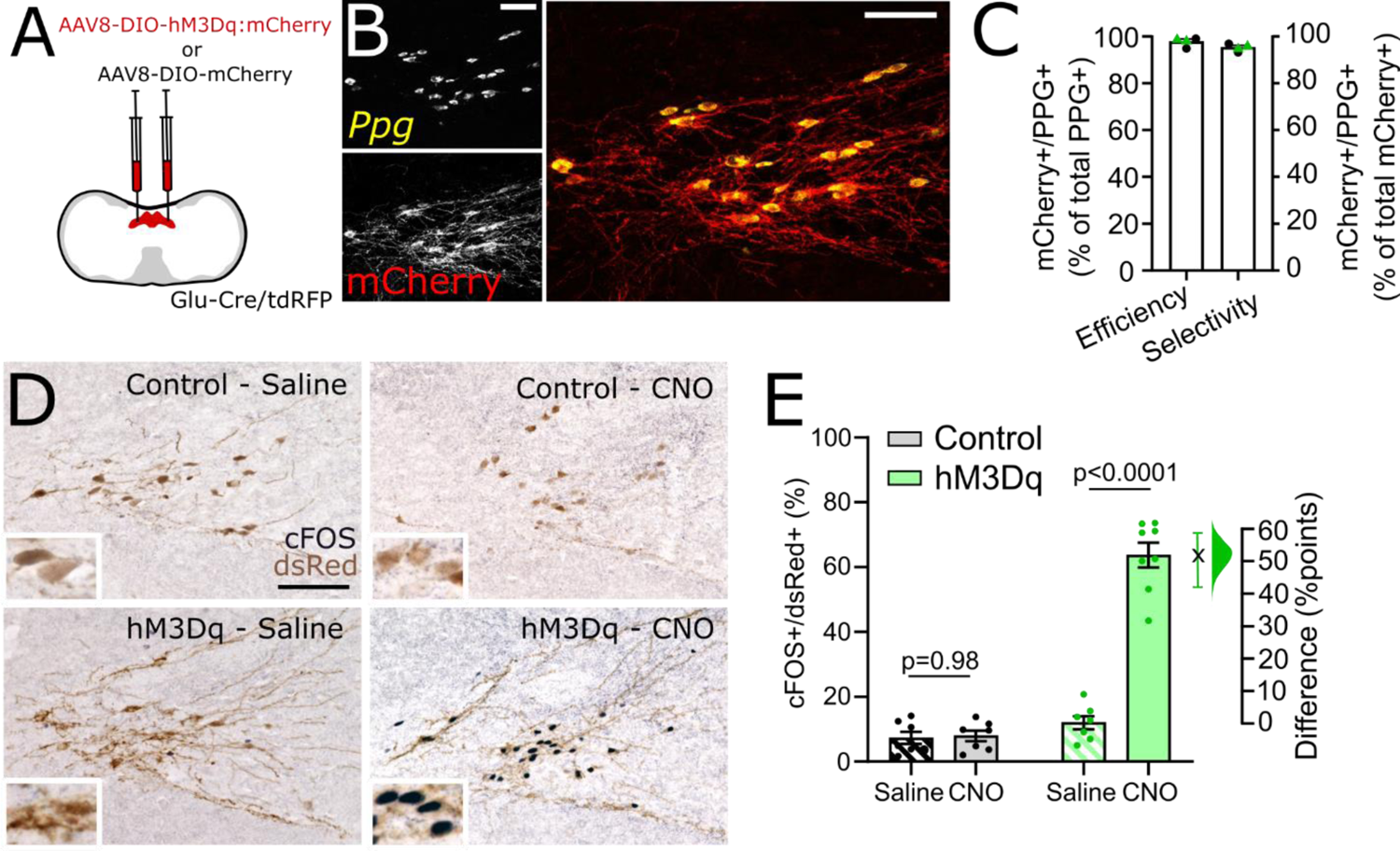
Selective and efficient chemogenetic activation of cNTS PPG neurons *in vivo.* A) Schematic of injection protocol. B) Representative images of RNAscope *in situ* hybridisation for *Ppg* mRNA (yellow) and immunolabelling for mCherry (to detect hM3Dq:mCherry, red) in tissue from one Glu-Cre/tdRFP mice injected with AAV8-DIO-hM3Dq:mCherry. Scale bars: 100µm. C) Percent of *Ppg-*expressing neurons in the cNTS also expressing hM3Dq:mCherry (efficiency), as well as percent of mCherry-expressing cNTS neurons also expressing *Ppg* (selectivity). Results from control mice (expressing mCherry only, n=2) are indicated with black circles, while results from hM3Dq-expressing mice (n=2) are indicated with green triangles. D) Representative images of immunohistochemical labelling for cFOS (black nuclear label) and dsRed (brown cytoplasmic label, detecting mCherry and tdRFP) in mice expressing mCherry only (control, top panels) or hM3Dq:mCherry (hM3Dq, bottom panels) injected with saline (2ml/kg, left panels) or CNO (2mg/kg, 2ml/kg; right panels). Scale bar: 100µm. E) Percent of mCherry-expressing cNTS neurons also labelled for cFOS in control (grey/black) and hM3Dq-expressing (green) mice injected with saline (2ml/kg, pattern) or CNO (2mg/kg, filled). Also shown is Gardner-Altman estimation plot showing the mean difference in activated neurons between saline and CNO-injected hM3Dq-expressing mice. Two-way drug x virus interaction: F(1,25)=93.78, p<0.0001.

When data from male and female mice were combined, chemogenetic activation of PPG neurons suppressed cumulative food intake during six hours after dark onset (Fig 2A), consistent with previous findings (Brierley et al., 2021; Gaykema et al., 2017; Holt, Richards, et al., 2019; Liu et al., 2017). This prolonged suppression in feeding was driven by females eating significantly less following chemogenetic activation of PPG neurons (Fig 2B; treatment x virus x time interaction: F(5, 70)=6.889, p<0.0001). In contrast, there was no significant three-way interaction in male mice [Fig 2C, F(5, 75)=0.7909, p=0.56]. However, at 1 hour after dark onset, feeding was suppressed to a similar degree in both sexes following chemogenetic activation of PPG neurons (Fig S2A, Fig 2D, effect size: −0.187g [95%CI: −0.278g, −0.109g]). The latency to initiate the first meal was increased by 25.5 mins ([95%CI: 12.3 mins, 49.3 mins], Fig 2E) following chemogenetic activation, with no effect of sex (Fig S2B). Furthermore, the size of that first meal was reduced by 0.086g ([95%CI: 0.039g, 0.137g], Fig 2F), with no effect of sex (Fig S2C). There was no effect of CNO on the number of meals over the first six hours or on the average time taken between meals (Fig S2D, E). Interestingly, PPG neuron activation reduced average meal size by 0.091g (95%CI: 0.138g, 0.054g) in female mice over the first 6h of the dark phase, whereas the effect in males was more transient (Fig 2G). Chemogenetic activation had no impact on bodyweight 24 hours after injection of CNO (Fig 2H, Fig S2F), and CNO did not impact bodyweight or any assessed food intake metric in virus control mice.

**Figure 2.**
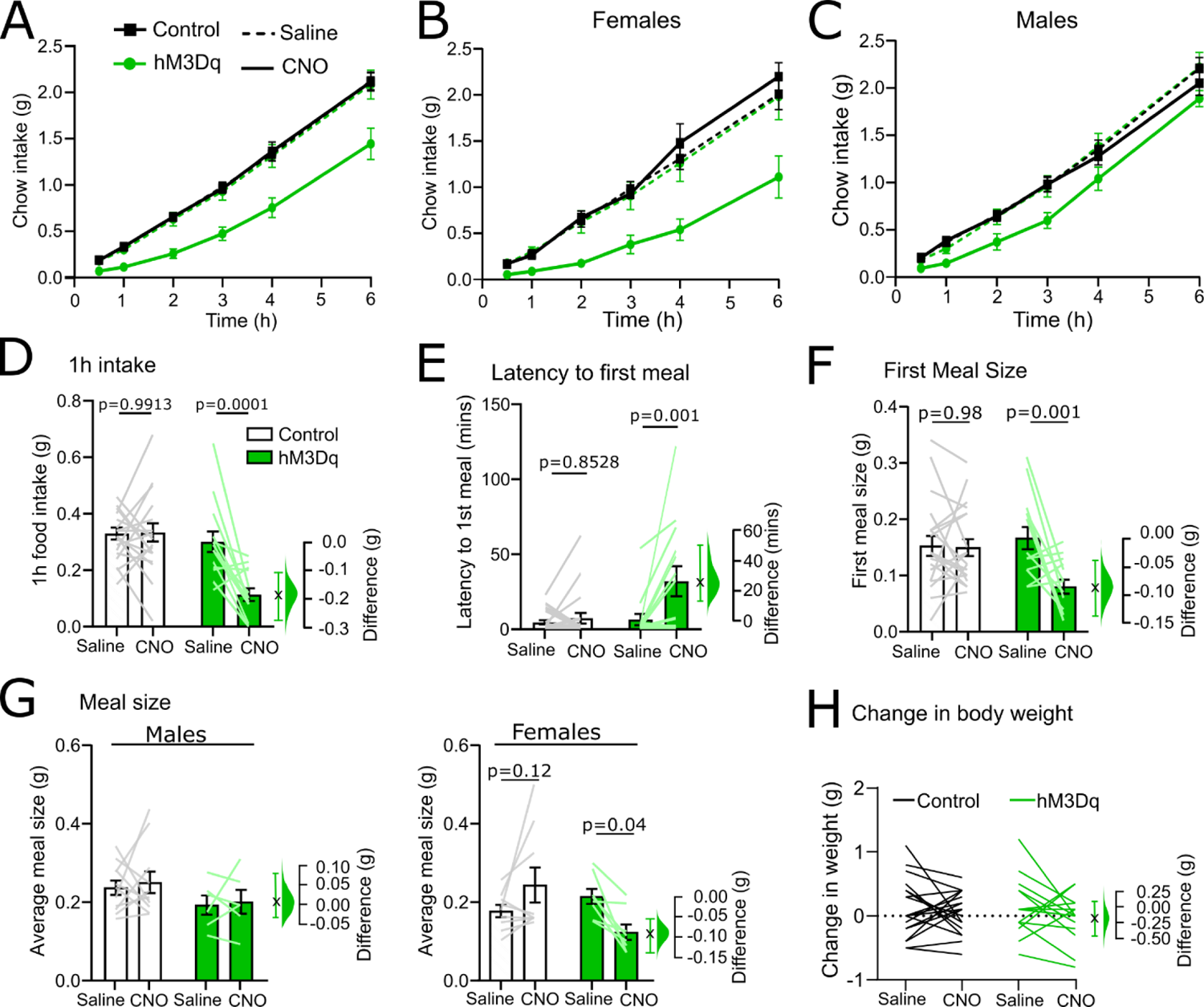
Meal pattern analysis of chow intake following chemogenetic activation of cNTS PPG neurons *in vivo.* A) Cumulative chow intake over the first six hours of the dark phase of male and female control (black squares) and hM3Dq-expressing mGlu-Cre/tdRFP mice (green circles) following i.p. injection of either saline (dashed lines, 2ml/kg) and CNO (solid lines, 2mg/kg, 2ml/kg). Three-way drug x time x virus interaction: F(5,110)=3.628, p=0.0045. B) Cumulative chow intake of females only. Three-way drug x time x virus interaction: F(5,70)=6.889, p<0.0001. Two-way drug x time (hM3Dq): F(5,35)=9.625, p<0.0001. Two-way drug x time (control): F(1.499, 10.49)=0.8945, p=0.4088. C) Cumulative chow intake of males only. Three-way drug x time x virus interaction: F(5,75)=0.7909, p=0.5595. Main effect of time [F(1.429, 21.43)=436.3, p<0.0001]. D) Chow intake by male and female mice one hour after dark onset [drug x virus: F(1, 32)=13.17; p=0.0010]. E) Latency of male and female mice to begin feeding [drug x virus: F(1, 32)=7.058, p=0.0122]. F) Size of the first meal in male and female mice [drug x virus: F(1, 32)=7.895; p=0.0084]. G) Average meal size over the first six hours grouped by sex in control (white bars) and hM3Dq-expressing mice (green bars) injected with saline (2ml/kg) or CNO (2mg/kg, 2ml/kg). Three-way drug x virus x sex interaction: F(1, 30)=4.882; p=0.0349; two-way drug x virus (females): F(1, 15)=11.11, p=0.0045; two-way drug x virus (males): F(1, 15)=0.01089, p=0.9183). H) No significant change in bodyweight of control (black) or hM3Dq-expressing mice (green) 24 hours after injection of saline (2ml/kg) or CNO (2mg/kg, 2ml/kg) [virus x drug: F(1, 32)=0.4636, p=0.5008]. The difference in the relevant outcome between saline and CNO-injected hM3Dq-expressing mice is shown in D-H using Gardner-Altman estimation plots.

### Activation of PPG neurons is sufficient to induce mild anxiety-like behaviour

Increased latency to begin feeding and short-term suppression in food intake can be indicative of negative affect, including anxiety-like states (Holt & Trapp, 2016). While a previous study failed to find any impact of chemogenetic activation of PPG neurons on anxiety-like behaviours in mice (Gaykema et al., 2017), results from several studies in rats have revealed anxiogenic effects of GLP1 receptor stimulation in multiple brain regions (Kinzig et al., 2003; López-Ferreras et al., 2020; Zheng et al., 2019). Given this apparent discrepancy in the rodent literature, we next investigated the extent to which chemogenetic activation of PPG neurons is sufficient to induce anxiety-like behaviour in mice using two behavioural assays. First, as in Gaykema et al. (2017), we analysed exploratory behaviour of mice placed into a novel open field, a validated test for anxiety-like behaviour that relies on rodents’ innate avoidance of open, exposed areas and their tendency to stay close to corners and edges (Seibenhener & Wooten, 2015; Walsh & Cummins, 1976). Only male mice were used in the study by Gaykema et al. (2017, personal communication). We found that female and male mice behaved differently in the open field (main effect of sex: p=0.03), with males exhibiting significantly higher variability in the time spent in the centre of the field (Fig 3C). In female mice, chemogenetic activation of PPG neurons significantly reduced the time spent in the centre of the open field by 71s (95%CI: 29.3s, 116s; Fig 3A,B), and reduced the total distance travelled by 29.5m (95%CI: 0.5m, 48.5m, Fig 3A,B). In male mice, chemogenetic activation had no effect on time spent in the centre of the open field or on total distance travelled (Fig 3C).

**Figure 3.**
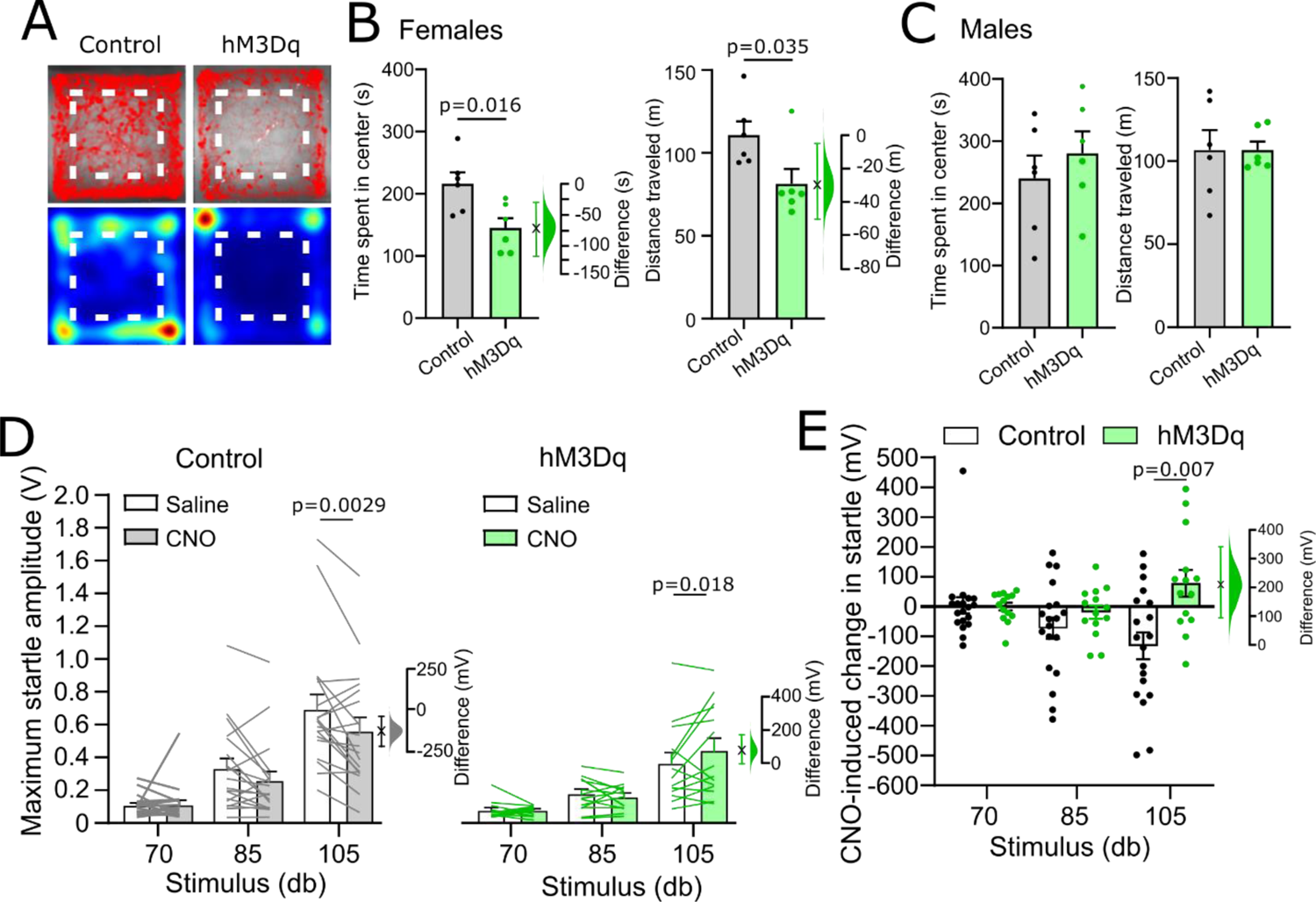
Chemogenetic activation of cNTS PPG neurons is sufficient to elicit moderate increases in anxiety-like behaviour. A) Traces and heatmaps from a representative control (left) and hM3Dq-expressing mouse (right) recorded in the open field following injection of CNO (2mg/kg, 2ml/kg). The centre of the arena is outlined using a dashed line. B-C) Quantification of time spent in the centre of the open field (left) and the total distance travelled (right) in female (B) and male (C) control (white bars) and hM3Dq-expressing mice (green bars). Student’s T-test: females, centre time: t=2.912, df=10; females, distance: t=2.438, df=10; males, centre time: t=0.7911, df=10; males, distance: t=0.003524, df=10. D) Maximum acoustic startle amplitude of control (grey/white) and hM3Dq-expressing (green) male and female mice (combined due to no sex difference [Fig S3]) following injection of saline (2ml/kg) or CNO (2mg/kg, 2ml/kg). Three-way drug x virus x db interaction: F(2, 62)=5.233; p=0.0079. Two-way drug x db interaction (control): F(2, 36)=3.435; p=0.0431. Two-way drug x db interaction (hM3Dq): F(2, 26)=3.904; p=0.0329. E) CNO-induced startle calculated as the difference in startle between injection of saline and CNO for each animal. Control mice: black circles, white bars; hM3Dq-expressing mice: green circles, green bars. Two-way virus x db interaction: F(2, 62)=5.233; p=0.0079. Also shown in B), D) and E) is the effect size using a Gardner-Altman estimation plot.

A second test investigated anxiety-like behaviour in male and female mice by assaying acoustic startle response magnitude, which does not depend on locomotion or exploratory behaviour. Another benefit of the acoustic startle test is that it can be conducted using a repeated-measures, within-subject design (Plappert et al., 2005). There was no significant difference between males and females in the startle response to three different noise (db) levels (main effect of sex control group: p=0.21, main effect of sex hM3Dq group: p=0.61). However, there was a significant three-way drug x virus x db interaction [F(2, 62)=5.233, p=0.0079, Fig 3D]. Follow-up analyses revealed a significant two-way drug x db interaction in both the control [F(2, 36)=3.435, p=0.043)] and hM3Dq group [F(2, 26)=3.904, p=0.033]. CNO (2mg/kg i.p.) reduced startle responses at the highest noise level (105 db) in control mice (effect size: −132mV with a 95%CI of [−40.04mV, −223.6mV]), consistent with a previous report in Long-Evans rats treated with CNO (MacLaren et al., 2016). This effect of CNO on acoustic startle appeared to be more pronounced in male mice, although the effect of sex was not significant (Fig S3). Conversely, in mice expressing hM3Dq, CNO injection (2mg/kg) *increased* startle amplitude responses to the 105 db noise by 78.1mV (95%CI: −2.31mV, 168mV). This differential CNO effect in control and hM3Dq mice became even more apparent when the difference between startle amplitude in the presence of saline vs. CNO was plotted: CNO-induced increases in startle amplitude within subjects was 210mV higher in hM3Dq mice compared to control mice (95%CI: 96.8mV, 337mV, Fig 3E).

### Monosynaptic input from the PVN drives PPG neuron activation in male mice

Hypophagia and anxiety-like behaviours are driven by neural circuits whose activity is modulated by acute and chronic stress (Holt, 2021; Holt & Rinaman, 2021; Ulrich-Lai & Herman, 2009; Zheng et al., 2019). Having revealed a potential role for PPG neurons not only in food intake but also in anxiety-like behaviour in mice, we next sought to identify neural pathways which may drive activation of PPG neurons in response to acute stress. Rabies-mediated monosynaptic tracing (Fig 4A, Fig S4A) confirmed our previous findings that multiple forebrain, midbrain, and hindbrain regions provide direct input to NTS PPG neurons (Holt, Pomeranz, et al., 2019), with particularly prominent afferent inputs arising from the PVN (Fig S4B). Prior to perfusion, a subset of these mice was exposed to 30mins restraint stress (Fig 4A). Compared to non-stressed controls, restraint stress increased the percentage of PPG-projecting PVN neurons expressing cFOS by 27.1 percentage points (95%CI: 16.1, 47.8 percentage points; Fig 4B,C), supporting a role for these PVN neurons in stress-induced activation of postsynaptic PPG neurons.

**Figure 4.**
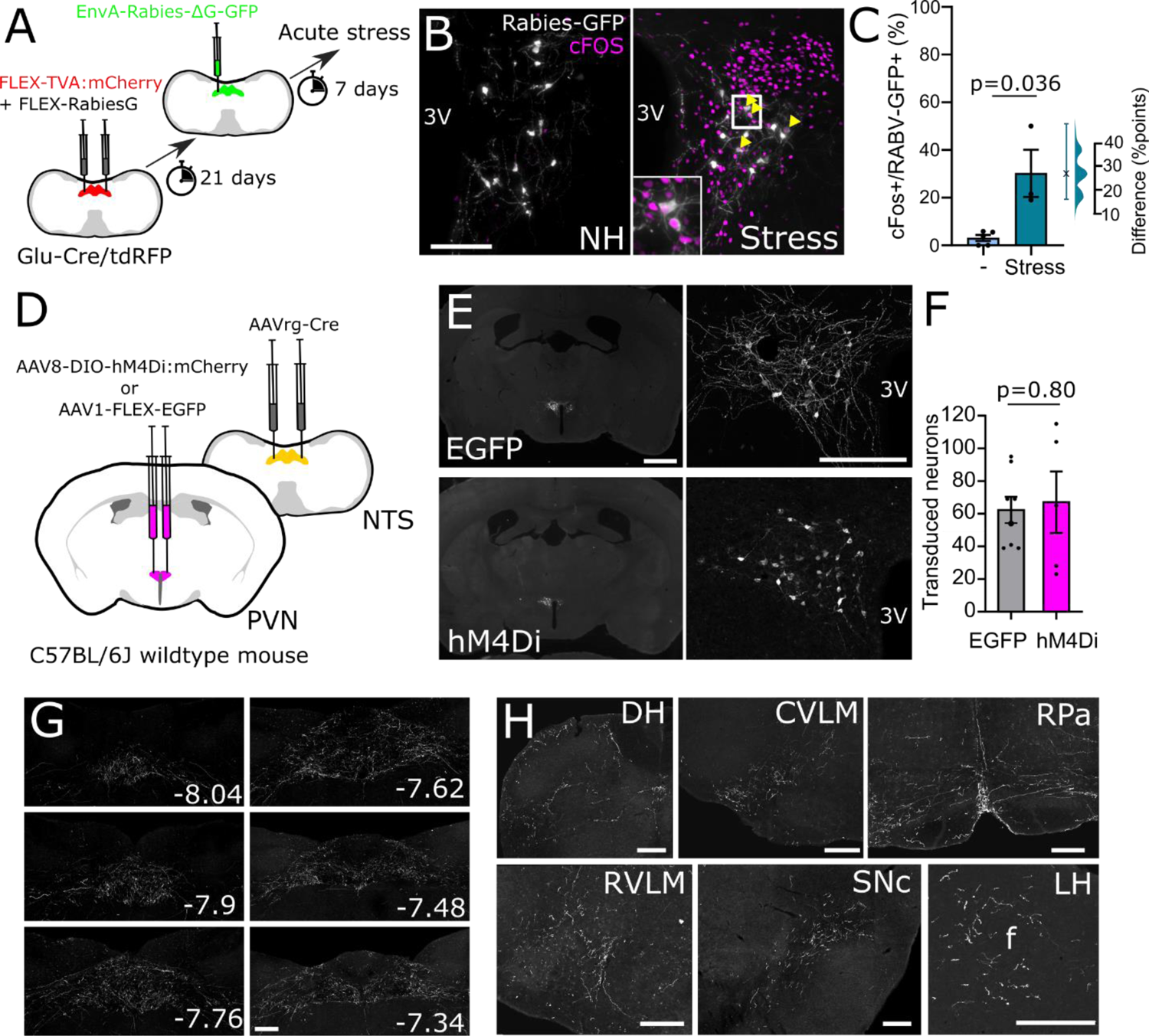
Identification and transduction of a stress-activated PVN→NTS circuit targeting PPG neurons. A) Monosynaptic input from the PVN to cNTS PPG neurons. B) GFP immunofluorescence (greyscale) in the PVN 7 days after unilateral microinjection of (EnvA)-RABV-ΔG-GFP targeted to the cNTS. cFOS immunoreactivity (magenta nuclei, pseudocolour) and RABV-GFP immunofluorescence (greyscale) is shown in a representative non-handled mouse (NH, left) and in a mouse perfused 90mins after the onset of 30mins restraint stress (Stress, right). Scale bar: 100µm. C) Calculated percentage of RABV-GFP labelled cells that were also cFOS-positive in nonhandled mice (-, n=5) vs mice exposed to 30mins restraint stress (n=3). Mann-Whitney U test: t=3.203, df=5. Also shown is the difference between EGFP and hM4Di-expressing mice in the percentage of RABV-GFP labelled cells activated to express cFOS using a Gardner-Altman estimation plot. D) Schematic illustrating the intersectional approach for targeting of NTS-projecting PVN neurons to induce Cre-dependent expression of EGFP or hM4Di. E) Representative images at low (left, scale bar: 1mm) and high magnification (right, scale bar: 200µm) of immunolabelling for EGFP (top) or hM4Di:mCherry (bottom) in NTS-projecting PVN neurons. 3V: third ventricle. F) Similar numbers of NTS-projecting PVN neurons transduced to express either EGFP or hM4Di:mCherry. G) Representative images of PVN-derived, EGFP-labelled axons throughout the cNTS in mice expressing EGFP in NTS-projecting PVN neurons. Scale bar: 100µm. Numbers indicate distance from bregma in mm. H) Additional axon collateral targets of NTS-projecting PVN neurons. DH: dorsal horn, CVLM: caudal ventrolateral medulla, RPa: raphe pallidus, RVLM: rostral ventrolateral medulla, SNc: substantia nigra pars compacta, LH: lateral hypothalamus. Scale bars: 100µm.

We next used an intersectional approach to acutely inhibit NTS-projecting PVN neurons in male wildtype mice, including (but not limited to) neurons that provide synaptic input to PPG neurons. For this, a retrogradely transported AAV (AAVrg-hSyn-Cre) was microinjected into the cNTS in male C57BL/6J mice to induce Cre expression in NTS-projecting neurons throughout the brain, combined with injection of AAV8-hSyn-DIO-hM4Di:mCherry or AAV1-CAG-FLEX-EGFP into the PVN to induce selective expression of hM4Di:mCherry or EGFP in NTS-projecting PVN neurons (Fig 4D,E). There was no difference in the number or distribution of neurons transduced with either virus (Fig 4F). In virus control mice, EGFP labelling filled the cytoplasm and processes of transduced PVN neurons, including their axon collaterals, revealing other brain regions that receive collateralized input from NTS-projecting PVN neurons. As expected, the NTS contained the densest accumulation of axonal GFP-immunolabelling, which extended through the rostrocaudal extent of the NTS (Fig 4G). Additional GFP-immunolabelled fibres representing the axon collaterals of NTS-projecting PVN neurons were seen in the dorsal horn of the spinal cord, the caudal ventrolateral medulla, the raphe pallidus, the rostral ventrolateral medulla, the substantia nigra, and the lateral hypothalamus (Fig 4H). No GFP-positive fibres were observed within telencephalon regions, within diencephalic regions rostral to the PVN, or within the median eminence (Fig S4C).

To determine the extent to which recruitment of NTS-projecting PVN neurons contributes to stress-induced activation of GLP1 neurons, EGFP and hM4Di-expressing mice were injected with CNO to inhibit NTS-projecting PVN neurons. Mice were then exposed to either 30mins restraint or 20mins in a novel environment, both of which are acutely stressful to rodents (Tsukiyama et al., 2011) (Fig 5A). There was no difference between stressors in their ability to activate cFOS, and so data were pooled for further analysis. As expected, CNO-induced chemogenetic inhibition attenuated stress-induced activation of hM4Di-expressing PVN neurons by 13.6 percentage points (95%CI: 7.37, 23.2 percentage points) compared to PVN activation in EGFP-expressing controls (Fig 5B,C). Further, acute inhibition of NTS-projecting PVN neurons was sufficient to reduce stress-induced activation of neurons within the cNTS by 19.4 counts/section (95%CI: 9.9, 29.4 counts/section, Fig 5D,E). The effect was even larger among GLP1-positive cNTS neurons, for which the proportion of GLP1 neurons expressing cFOS was reduced by 26.3 percentage points (95%CI: 13.7, 39.6 percentage points, Fig 5F,G). These results provide evidence that NTS-projecting PVN neurons contribute to stress-induced activation of PPG/GLP1 and other NTS neurons.

**Figure 5.**
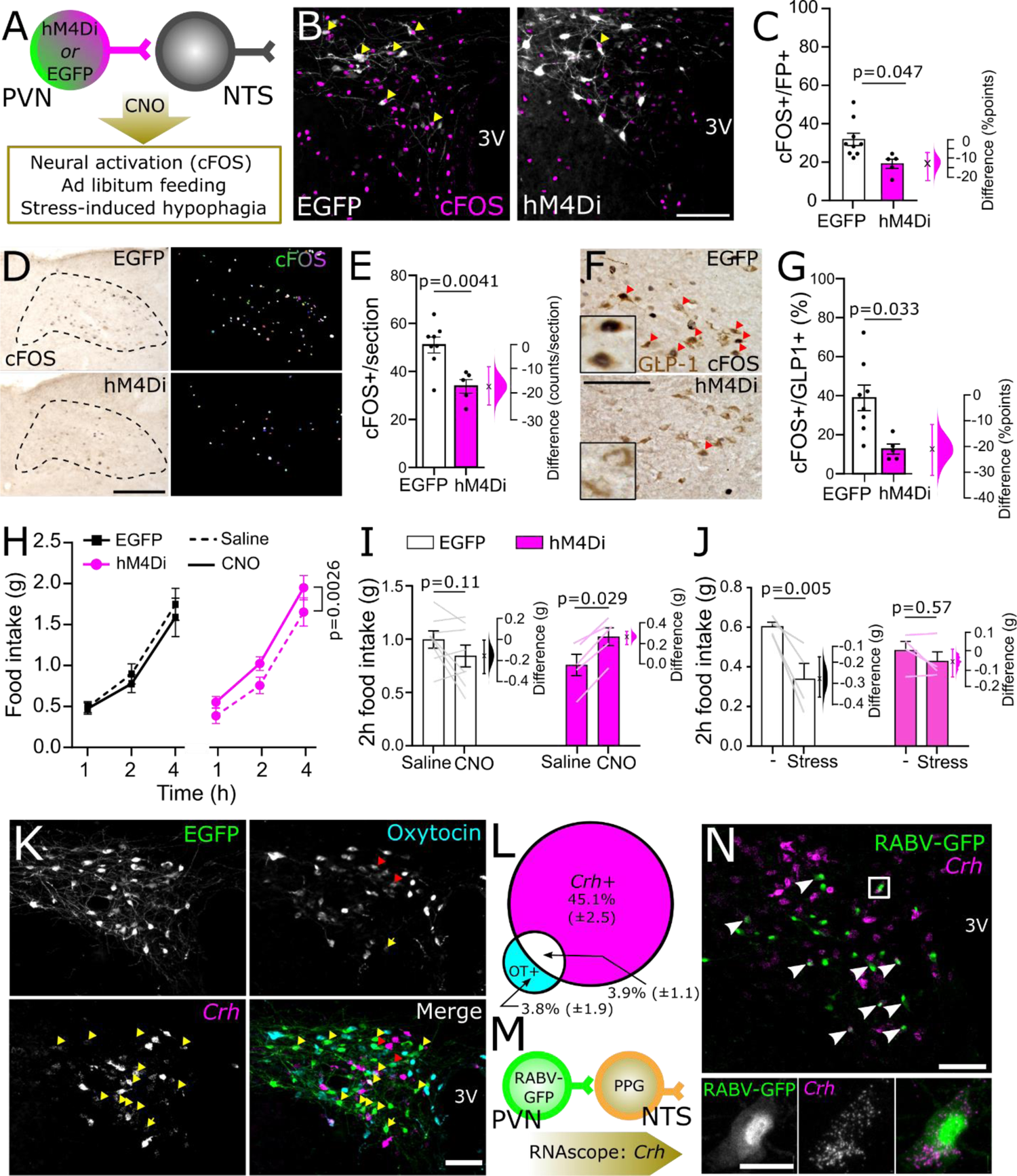
Chemogenetic inhibition of NTS-projecting PVN neurons. A) Diagram of the experimental paradigm for chemogenetic inhibition of NTS-projecting PVN neurons. B) cFOS immunoreactivity (magenta nuclei, pseudocolour) and EGFP or mCherry immunofluorescence (greyscale) labelling in the PVN of a representative control mouse (left, EGFP) and a representative hM4Di-expressing mouse (right, hM4Di) after i.p. injection of CNO (2mg/kg, 2ml/kg) followed by 30mins restraint stress. 3V: third ventricle. Scale bar: 100µm. C) Calculated percentage of EGFP-vs. hM4Di:mCherry-expressing (fluorescent protein, FP+) cells that were also cFOS-positive in mice after i.p. injection of CNO (2mg/kg, 2ml/kg) followed by exposure to an acute stressor (novel environment or restraint stress). Unpaired T-test: t=2.498, df=6. D) Representative images of cFOS-immunoreactivity in the cNTS of mice expressing EGFP (top) or hM4Di (bottom) in NTS-projecting PVN neurons. Counted cFOS-immunoreactive cells indicated with rainbow-coloured mask to the right. All mice were injected with CNO (2mg/kg, 2ml/kg) and then exposed to novel environment or restraint stress. Scale bar: 100µm. E) Counts of cFOS-immunoreactive nuclei in the cNTS in mice expressing EGFP or hM4Di in NTS-projecting PVN neurons and injected with CNO (2mg/kg, 2ml/kg) prior to stress exposure. Unpaired T-test: t=3.603, df=11. F) cFOS- and GLP1-immunoreactivity in the cNTS of mice expressing EGFP (top) or hM4Di (bottom) in NTS-projecting PVN neurons and injected with CNO (2mg/kg, 2ml/kg) prior to stress exposure. Scale bar: 100µm. G) Calculated percentage of GLP1-immunoreactive cells in the cNTS that were also cFOS-IR. Unpaired T-test: t=2.766, df=6. H) Cumulative chow intake over four hours after dark onset in mice expressing EGFP or hM4Di in NTS-projecting PVN neurons after injection with saline (2ml/kg; dashed lines) or CNO (2mg/kg, 2ml/kg; solid lines). I) Chow intake over the first two hours of dark phase in mice expressing EGFP or hM4Di in NTS-projecting PVN neurons after injection with saline (2ml/kg) or CNO (2mg/kg, 2ml/kg). J) Chow intake during the first two hours of dark onset of mice expressing EGFP or hM4Di in NTS-projecting PVN neurons and injected with CNO (2mg/kg, 2ml/kg) prior to 30mins restraint stress. Virus x stress interaction: F(1, 6) = 7.805, p=0.0314. K) Representative images of RNAscope *in situ* hybridisation for *Crh* (magenta in merged image) combined with immunolabelling for GFP (green in merged image) and oxytocin (cyan in merged image) in the PVN of mice expressing GFP in NTS-projecting PVN neurons. Scale bar: 100µm. L) Percentage of NTS-projecting PVN neurons expressing *Crh* and/or oxytocin (OT+). N=3 mice, 2 sections containing the PVN from each mouse. M) Diagram illustrating RNAscope *in situ* labelling for *Crh* in cells with direct projections to NTS PPG neurons (RABV-GFP labelled). N) RNAscope *in situ* hybridisation for *Crh* in PPG-projecting PVN neurons (RABV-GFP labelled). Scale bar top panel: 100µm. Bottom panel: higher magnification image of inset in top panel. Scale bar: 20µm. Gardner-Altman estimation plots in panels C, E, G, I, and J display estimated effect sizes for each assessed parameter.

### NTS-projecting PVN neurons suppress feeding and drive stress-induced hypophagia

Chemogenetic inhibition of PVN neurons that project to the NTS was by itself sufficient to increase chow intake over the first four hours of dark onset (Fig 5H, main effect of drug in hM4Di group: p=0.0026). Two hours into the dark cycle, chow intake was increased by 0.266g [95%CI: 0.19, 0.316] in hM4Di-expressing mice injected with CNO compared to saline (Fig 5I). Conversely, in virus control mice, CNO tended to *suppress* food intake compared to intake after saline injection; however, the hypophagic effect in control mice was small and did not reach statistical significance (effect size: 0.154g [95%CI: 0.324, 0.00625]).

Since we previously found that NTS-projecting PVN neurons are activated in response to acute stress (Holt, Pomeranz, et al., 2019), we next tested whether chemogenetic inhibition of the PVN→NTS pathway was sufficient to reduce or block restraint stress-induced hypophagia. There was a large and statistically significant effect of restraint stress to reduce subsequent food intake in control mice, which consumed 0.265 g less over two hours following the stressed vs. non-stressed condition (Fig 5J; 95%CI: 0.155, 0.36g). In contrast, restraint stress-induced hypophagia was significantly attenuated in mice with acute inhibition of NTS-projecting PVN neurons, with mice eating only marginally less during the two-hour period after restraint stress vs. the non-stressed condition (effect size: 0.055 [95%CI: −0.01, 0.135g]).

To determine the neurochemical phenotype of NTS-projecting PVN neurons, RNAscope *in situ* hybridisation was performed to localise *Crh* mRNA expression combined with immunolabeling for oxytocin (OT) and viral reporter. Nearly half of identified NTS-projecting PVN neurons expressed *Crh*, but only a small number were OT-positive, and approximately half of the OT-positive projection neurons also expressed *Crh* mRNA (Fig 5K,L). Finally, to determine whether PVN^CRH^→NTS projection neurons are presynaptic to GLP1 neurons, we used RNAscope to localize *Crh* mRNA expression in tissue sections from one mouse with monosynaptic retrograde rabies virus labelling of PVN neurons that synapse onto NTS PPG neurons (Fig 5M,N). Of 59 retrogradely labelled PPG-projecting PVN neurons, the majority (37 neurons, 62.7%) expressed *Crh*.

## Discussion

Here we report that GLP1/PPG neurons have the ability to elicit anxiety-like behaviours in mice in a sex-dependent manner and that their activation following acute stress relies on a CRH-rich input from the PVN, which contributes to stress-induced suppression in feeding. Our finding that PPG neural activation suppresses feeding partly by decreasing the size of the first meal (with no impact on meal frequency) is consistent with acceleration of satiation, as reported previously by Brierley et al. (2021). However, an increase in the latency to begin feeding together with a decrease in first meal size can also be indicative of behavioural suppression, perhaps associated with negative emotional state (Calvez et al., 2011). Indeed, in addition to a well-established hypophagic role of central GLP1, solid evidence supports an anxiogenic role in rats, in which anxiety-like behaviours increase upon GLP1 receptor stimulation in the central amygdala (Kinzig et al., 2003) or supramammillary nucleus (López-Ferreras et al., 2020), and decrease following GLP1 receptor knockdown in the bed nucleus of the stria terminalis (Zheng et al., 2019). Our finding that PPG neural activation in female mice increases anxiety-like behaviour in the open field assay adds to the evidence for an anxiogenic role for GLP1/PPG neurons. A previous study failed to observe any anxiogenic effects of chemogenetic activation of PPG neurons when mice were tested in the elevated plus maze or open field (Gaykema et al., 2017). However, that study only tested male mice (Ronald Gaykema, personal communication) and only used behavioural assays dependent on exploratory behaviour, which is reduced by PPG neuron activation (Gaykema et al., 2017). Our new findings using the acoustic startle test indicate that both male and female mice respond to PPG neuron activation with an increased startle response indicative of increased arousal and/or vigilance, both of which are recognized components of anxiety-like behaviour in mice, rats, and humans (Bailey & Crawley, 2009).

### Potential sex differences

Previous studies, including our own, have demonstrated robust feeding suppression in mice following optogenetic or chemogenetic activation of PPG neurons. However, these studies reported data collected only in males (Gaykema et al., 2017; Holt et al., 2020; Liu et al., 2017), or were underpowered to reveal potential sex differences (Brierley et al., 2021; Cheng et al., 2020; Holt, Richards, et al., 2019). Here we document that after chemogenetic activation of PPG neurons, female mice displayed hypophagia for 6 hours, whereas the effect in males was more transient. In both sexes, food intake and body weight normalized within 24 hours after CNO treatment. Previous studies reported variable hypophagia effect durations in mixed-sex groups of mice after acute chemogenetic activation of PPG neurons, with results indicating either no persistent effect on 24h food intake and body weight (Holt, Richards, et al., 2019, this report) or a sustained effect on cumulative food intake and body weight lasting up to 48 hours after a single injection of CNO (Brierley et al., 2021). The reason for this discrepancy is unclear but may be due to differences in functional expression of the hM3Dq construct, including whether the population of PPG neurons in the adjacent intermediate reticular nucleus is transduced, and resulting variability in the level of activation.

Based on the evidence presented here and by Gaykema et al. (2017), we propose that PPG neural activation contributes to hypophagia and other behavioural indices of anxiety-like behaviour in a manner that is partially sex- and assay-dependent. In this regard, sex differences in other behavioural responses to GLP1 receptor stimulation and inhibition have been reported in rats (López-Ferreras et al., 2020). Collectively, these findings highlight the importance of including experimental subjects of both sexes in a variety of assays and support the hypothesis that the central GLP1 system modulates behaviour in a sex-dependent manner.

### Functional and anatomical evidence for a PVN-NTS pathway inhibiting feeding in response to stress

A recent study reported that hindbrain 5-HT contributes to stress-induced activation of GLP1 neurons in rats (Leon et al., 2021). Here we show that NTS-projecting PVN neurons are another necessary source of input for the ability of restraint stress to activate GLP1/PPG and other cNTS neurons in male wildtype mice. Our new finding is reminiscent of the demonstrated contribution of descending PVN axonal projections in stress-induced recruitment of noradrenergic NTS neurons in rats (Dayas et al., 2004; Li et al., 1996). Importantly, the axon collaterals of NTS-projecting PVN neurons targeted additional brain regions that may contribute to activation of PPG neurons. In this regard, there is evidence that PVN projections to the mesencephalic and pontine periaqueductal gray and parabrachial nuclei contribute to the feeding-suppressive effects of PVN stimulation (Stachniak et al., 2014). However, we observed very few labelled axon collaterals in those two regions arising from NTS-projecting PVN neurons. Considering our finding that approximately half of NTS-projecting PVN neurons express *Crh* mRNA, consistent with previous reports in rats (Ruyle et al., 2018; Sawchenko, 1987), it is possible that chemogenetic inhibition of NTS-projecting PVN neurons also attenuated ACTH and corticosterone release following acute stress. We did not measure plasma levels of these stress hormones, but the lack of axon collaterals in the median eminence originating from NTS-projecting PVN neurons is consistent with other evidence that neuroendocrine PVN neurons comprise a population that is distinct from brainstem-projecting PVN neurons (Gasparini et al., 2020; Stern, 2015; Swanson & Sawchenko, 1980). The existence of a monosynaptic circuit from the PVN to cNTS PPG neurons shown here and previously (Holt, Pomeranz, et al., 2019), coupled with evidence that PPG-projecting PVN neurons are activated after acute stress, support the view that a direct pathway from PVN to NTS contributes to stress-induced activation of GLP1/PPG neurons.

In addition to attenuating stress-induced hypophagia, acute inhibition of NTS-projecting PVN neurons increased dark-onset food intake in non-stressed male mice. This result was surprising, given previous reports (Maejima et al., 2019; Stachniak et al., 2014) that chemogenetic inhibition of PVN-derived axon terminals within the NTS failed to increase feeding in *ad libitum*-fed mice in the early hours of the light phase. The reason for this discrepancy in results between our study and the previous one(s) is unclear and should be addressed in future studies. The present examination of NTS-projecting PVN neurons was restricted to male mice due to limited availability of experimental animals during the COVID-19 pandemic. Future studies should explore the role of NTS-projecting PVN neurons in females, and should address whether this descending neural pathway contributes to other stress-related behaviours.

## Conclusions

In rodents, acute stressors increase vigilance and arousal, suppress food intake, and reduce exploratory and approach behaviours through engagement of complex neural circuits. Our findings highlight a hypothalamic-brainstem pathway from CRH neurons in the PVN to GLP1/PPG neurons in the cNTS in the modulation of behavioural responses to stress. We demonstrate that chemogenetic activation of GLP1-producing PPG neurons in the cNTS of mice suppresses food intake and increases anxiety-like responses in a partially sex-dependent manner. These findings along with previously published studies using rats and mice support the view that GLP1/PPG neurons are an integral component of neural circuits that modulate physiological and behavioural responses to acute stress, including increased heart rate (Ghosal et al., 2017; Holt et al., 2020), reduced feeding (Holt, Richards, et al., 2019; this report), and increased anxiety-like behaviour (this report, López-Ferreras et al., 2020; Zheng et al., 2019). We further demonstrate that PPG-projecting PVN neurons are activated by restraint stress, and that NTS-projecting PVN neurons are necessary for the ability of restraint stress to activate PPG and other NTS neurons in mice. Finally, we discovered that chemogenetic inhibition of NTS-projecting PVN neurons, approximately half of which express the stress neuropeptide CRH, increases food intake under baseline conditions and attenuates stress-induced hypophagia. These novel findings highlight this descending pathway within a broader neural system that orchestrates behavioural stress responses.

---

**Fig S1 Relating to Fig 1.**
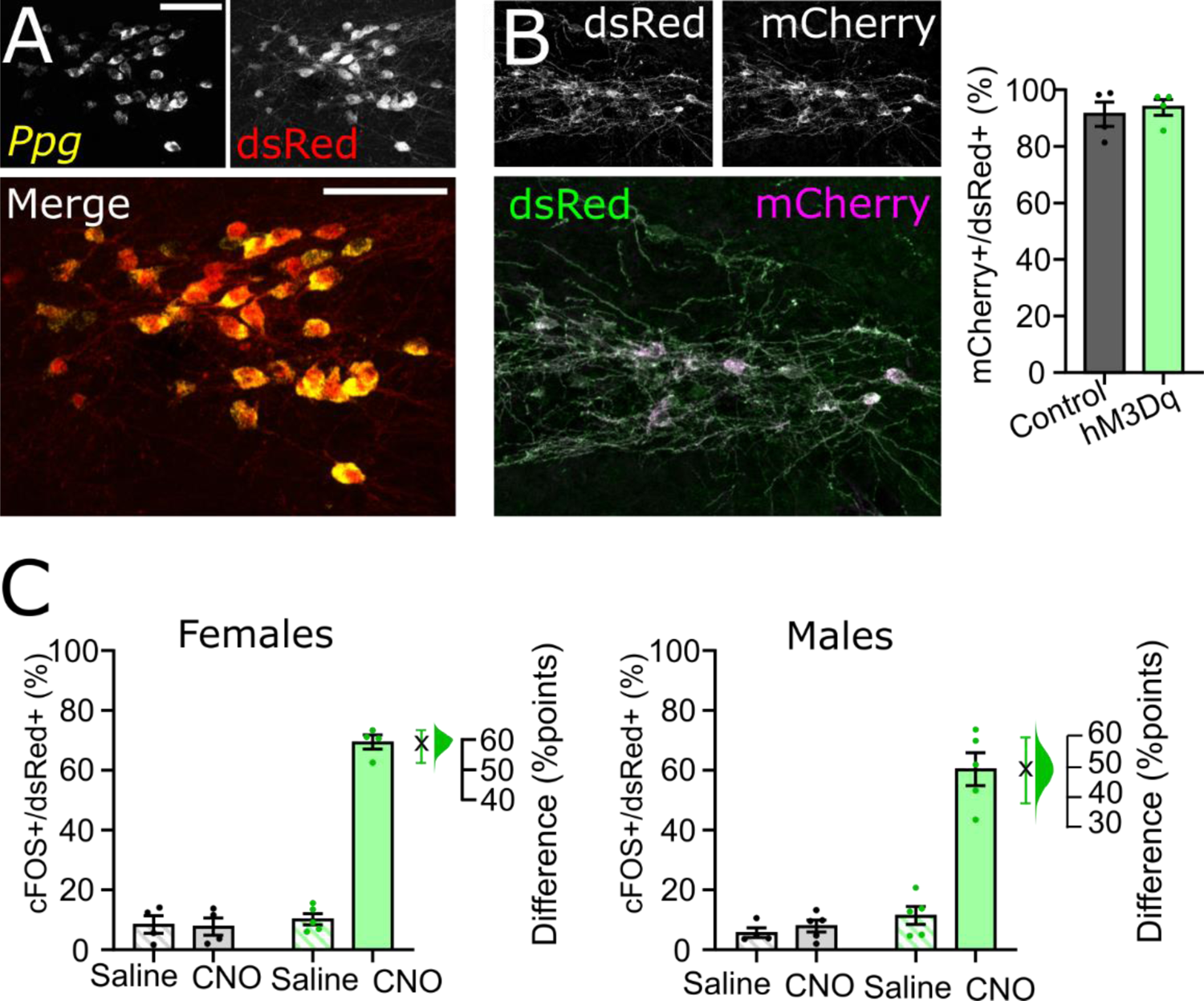
A) Immunolabelling for tdRFP (dsRed, red) and RNAscope *in situ* hybridisation for *Ppg* mRNA (yellow) in the cNTS of naïve Glu-Cre/tdRFP transgenic mice showing robust *Ppg* expression in tdRFP-positive cells with no discernible expression in non-*Ppg* cells in the cNTS. Scale bars: 100µm. B) immunolabelling for mCherry (detecting hM4Di:mCherry only, magenta) and dsRed (detecting both hM4Di:mCherry *and* tdRFP, green) in Glu-Cre/tdRFP mice injected with AAV8-hSyn-DIO-hM4Di:mCherry. Student’s T-test: p=0.66. C) Data from Fig 1E grouped by sex. Percent of mCherry-expressing cNTS neurons also labelled for cFOS in male and female control mice (grey/black) and hM3Dq-expressing mice (green) injected with saline (2ml/kg, striped) or CNO (2mg/kg, solid) expressing mCherry only or hM3Dq:mCherry (hM3Dq) after being injected with saline (2ml/kg) or CNO (2mg/kg, 2ml/kg). Gardner-Altman estimation plots display the estimated effect sizes. There was no three-way interaction of virus x drug x sex [F(1, 28)=2.151; p=0.1537].

**Fig S2 Relating to Fig 2.**
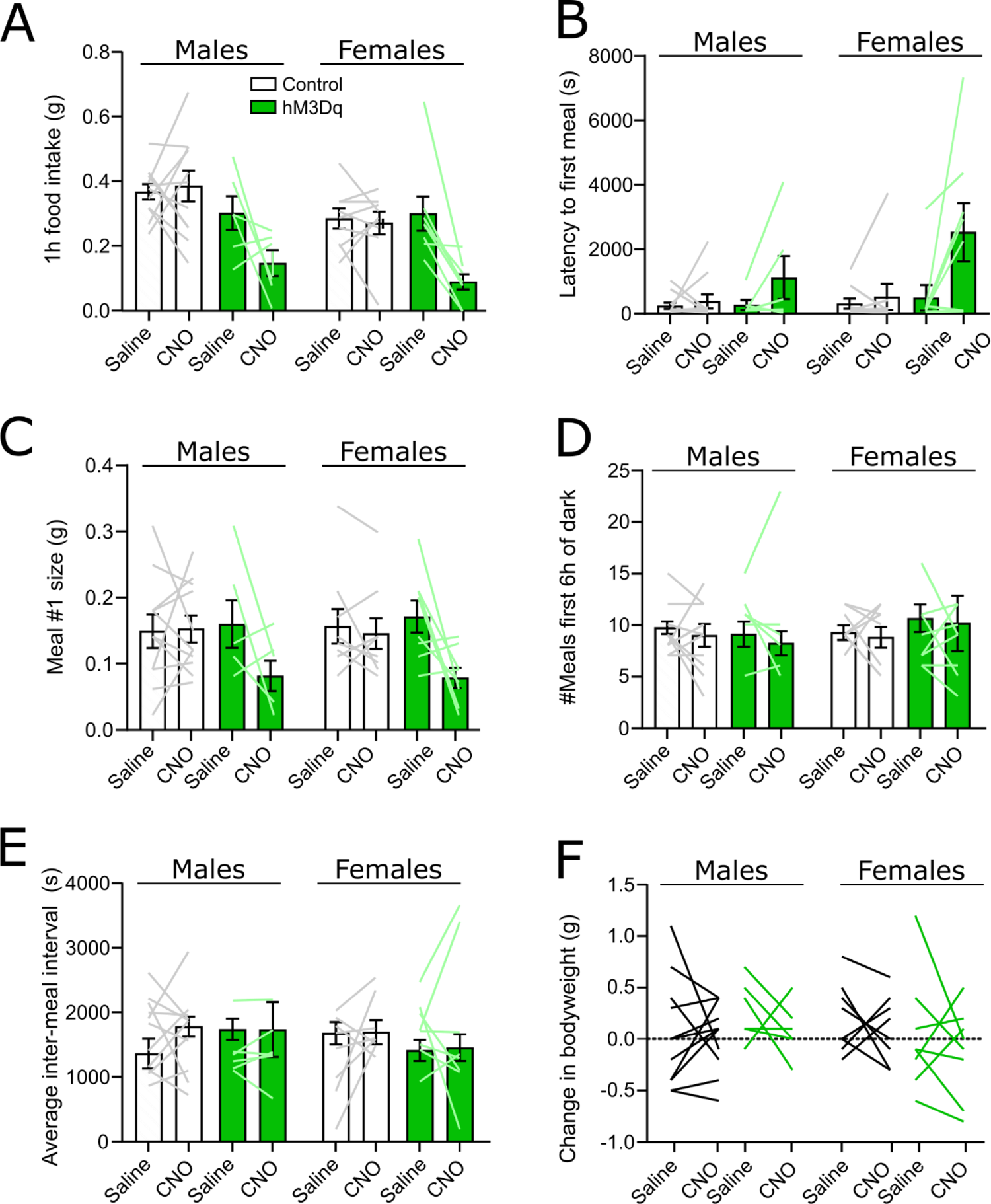
Meal pattern data grouped by sex in control mice (black) and hM3Dq-expressing mice (green) following injection of saline (2ml/kg) or CNO (2mg/kg, 2ml/kg). A) One hour food intake; no three-way drug x sex x virus interaction [F(1, 30)=0.05197; p=0.8212]. B) Latency to first meal; no three-way drug x sex x virus interaction [F(1, 30)=0.05197; p=0.8212]. C) First meal size; no three-way drug x sex x virus interaction [F(1, 30)=8.890e-005; p=0.9925]. D) Number of meals over the first six hours of the dark phase; no three-way drug x sex x virus interaction: F(1,29)=0.00079, p=0.977. E) Average inter-meal interval over the first six hours of the dark phase; no three-way drug x sex x virus interaction: F(1,29)=0.6371, p=0.4312. F) Change in body weight 24 hours after CNO or saline injection; no three-way drug x sex x virus interaction: F(1,29)=01381, p=0.7129.

**Fig S3 Relating to Fig 3.**
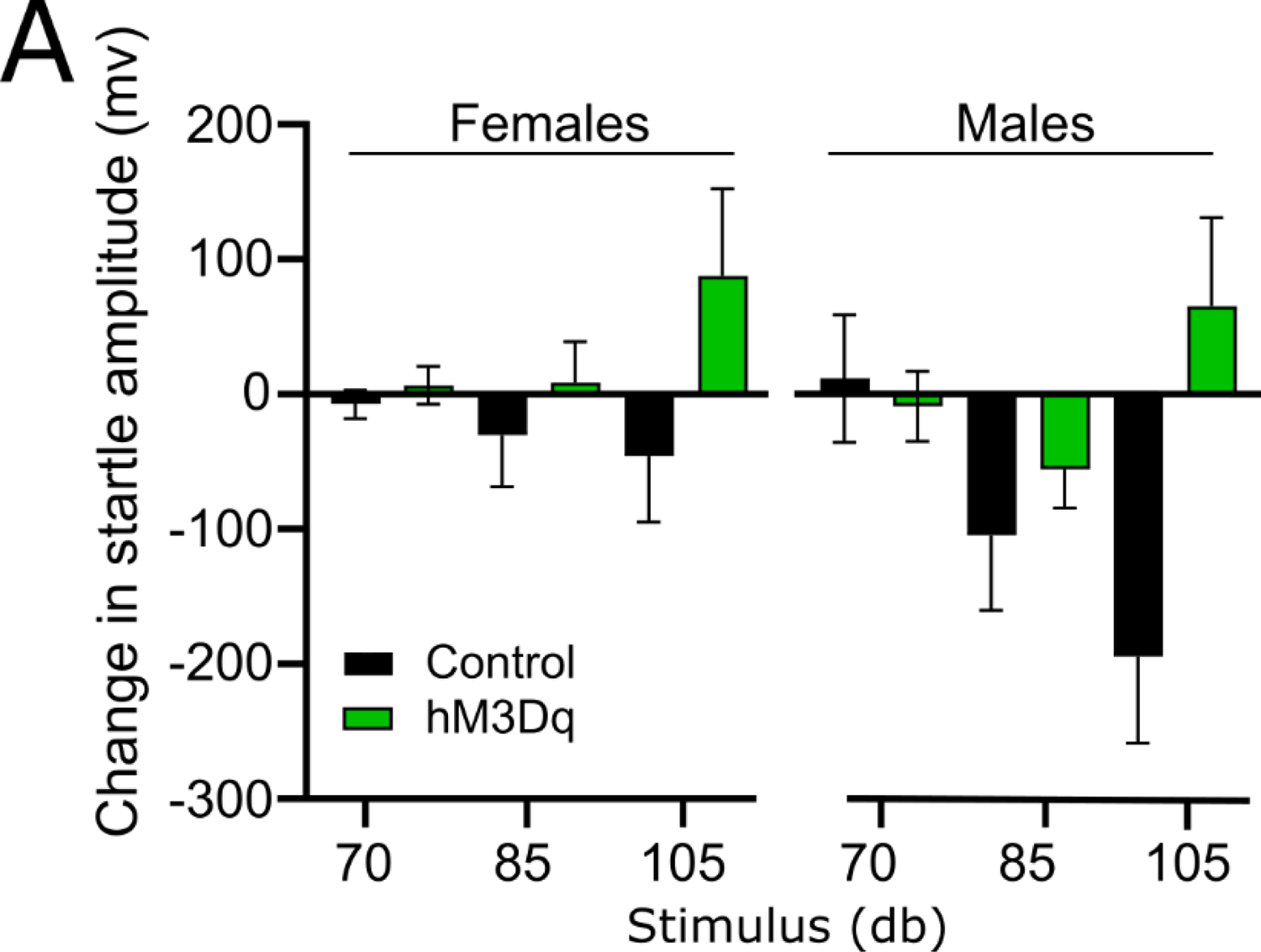
Acoustic startle results in female and male mice. A) Treatment effects on acoustic startle calculated as the within-subjects difference in startle amplitude in mice after injection of saline vs. CNO. No three-way drug x sex x virus interaction [F(2,58=0.721, p=0.491].

**Fig S4. Relating to Fig 4.**
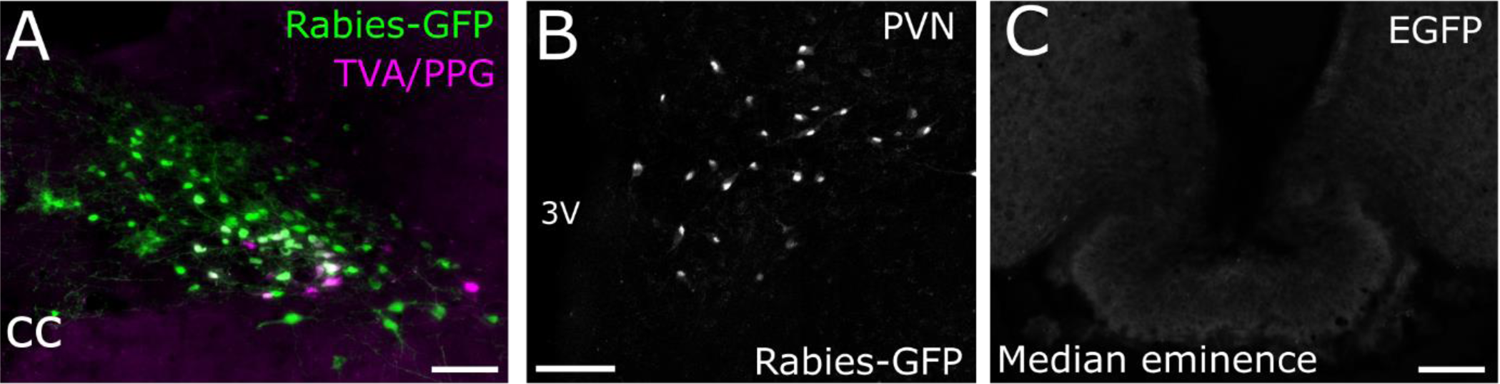
A) Representative image of RABV-GFP labelling (green) and dsRed-positive cells (magenta) expressing TVA receptor and/or tdRFP within the NTS. Scale bar: 100µm. B) RABV-GFP labelling in the PVN following monosynaptic tracing from PPG neurons. Scale bar: 100µm. C) No detectable axon collaterals within the median eminence arising from NTS-projecting PVN neurons. Scale bars: 100µm.

